# Preprocessing, normalization and integration of the Illumina HumanMethylationEPIC array

**DOI:** 10.1101/065490

**Authors:** Jean-Philippe Fortin, Timothy J. Triche, Kasper D. Hansen

**Affiliations:** Department of Biostatistics, Johns Hopkins Bloomberg School of Public Health; Jane Anne Nohl Division of Hematology, Keck School of Medicine of USC; McKusick-Nathans Institute of Genetic Medicine, Johns Hopkins School of Medicine

## Abstract

The *minfi* package is widely used for analyzing Illumina DNA methylation array data. Here we describe modifications to the *minfi* package required to support the HumanMethylationEPIC (”EPIC”) array from Illumina. We discuss methods for the joint analysis and normalization of data from the HumanMethylation450 (”450k”) and EPIC platforms. We also introduce the single-sample Noob (*ssNoob*) method, a normalization procedure suitable for incremental preprocessing of individual Human-Methylation arrays. Our results recommend the *ssNoob* method when integrating data from multiple generations of Infinium methylation arrays. Finally, we show how to use reference 450k datasets to estimate cell type composition of samples on EPIC arrays. The cumulative effect of these updates is to ensure that *minfi* provides the tools to best integrate existing and forthcoming Illumina methylation array data.

## Introduction

The IlluminaHumanMethylation450 (“450k”) array is a widely used platform for assaying DNA methylation in a larger number of samples using bisulfite conversion followed by array hybridization (Bibikova et al., 2011), and has been the platform of choice for epigenome-wide association studies and large scale cancer projects. In 2015, Illumina released their next generation methylation array, the HumanMethylationEPIC (“EPIC”) array (Moran, Arribas, and Esteller, 2016), with almost twice the number of CpG loci. This increased resolution, coupled with greatly expanded coverage of regulatory elements, makes the EPIC array an attractive platform for large-scale profiling of DNA methylation.

The minfi package in R/Bioconductor (Gentleman et al., 2004; Huber et al., 2015) is a widely used software package for analyzing data from the Illumina HumanMethylation450 array (Aryee et al., 2014). In addition to the analysis methods provided in the package, it exposes a flexible framework for handling DNA methylation data. Because of this, minfi is used as the data backend for many other software packages, including ChAMP (Morris et al., 2014), DMRcate (Peters et al., 2015), quantro (Hicks and Irizarry, 2015), missMethyl (Phipson, Maksimovic, and Oshlack, 2016), shinyMethyl (Fortin, Fertig, and Hansen, 2014), MethylAid (Iterson et al., 2014), ENmix (Xu et al., 2016), ELMER (Yao et al., 2015), conumee (Hovestadt and Zapatka, 2016), CopyNumber450k (Papillon-Cavanagh, Fortin, and De Jay, 2013), funtooNorm (Oros Klein et al., 2016) and Multi-DataSet (Ruiz-Arenas, Hernandez-Ferrer, and Gonzalez, 2016).

## Results

### Extending the minfi package to handle EPIC arrays

Here we report the extension of the minfi package to handle the recently released HumanMethylationEPIC DNA methylation microarray from Illumina, also referred to as the “EPIC” or “850k” array. Simultaneously, we extend the interactive visualization tool shinyMethyl (Fortin, Fertig, and Hansen, 2014), used for quality control assessment of 450k datasets, to add support for the EPIC array.

The launch of the EPIC array has been marred by some “technical” difficulties, including (1) issues with software settings in the scanner and (2) issues with the released annotation files describing the array design. For (1), early scanner settings distributed by Illumina incorrectly masked certain probes, resulting in an IDAT file with fewer registered probes (1,052,641 probes / 866,836 methylation loci in current IDAT files, versus 1,032,279 probes / 855,184 methylation loci in early access files, representing a difference of 11,652 methylation loci). This can be addressed by rescanning the physical array with updated scanner settings, which is sometimes impossible. The presence of multiple valid IDAT files for the same array design has led to the creation of a check for this situation in the parsing functions in minfi. For (2), Illumina has released multiple different versions of their “manifest” file which serves the dual purpose of describing the array design and annotation of the measured methylation loci. The different versions of the manifest file differ in a small number of probes. These errors do not reflect errors in the manufacturing process, but rather errors in the annotation released to outside researchers.

Because of these issues, it is critical (at the time of writing) to keep minfi and its associated annotation packages updated to their latest versions. To keep the associated annotation packages current, it may be necessary to re-read the original IDAT files with newer versions of minfi; the resulting R object will be linked to the latest annotation package.

For methods development and software testing, we have bundled 3 replicates of the GM12878 cell line assayed on the EPIC chip and released by Illumina as a demo dataset into an experimental package named minfiDataEPIC, available from Bioconductor. These 3 replicates are used below in a number of analyses.

Additionally, we have developed similar annotation and data packages for the older Illumina HumanMethylation27 (”27k”) array; this required extending internal functions in minfi to work with array designs having only one probe type. The minfiData27k package bundles 20 kidney samples (tumor and normal) assayed on the 27k array from the TCGA Kidney Renal Papillary Cell Carcinoma (KIRP) study (Cancer Genome Atlas Research Network, 2013).

The different array types along with the currently available preprocessing functions are listed in Table 1.

**Table 1.** Summary of the Illumina Methylation Arrays.

| Array | N. CpGs | Type I | Type II | Preprocessing functions in minfi | Data package |
| --- | --- | --- | --- | --- | --- |
| 27k | 27,578 | 27,578 | 0 | preprocessRaw, preprocessNoob, preprocessQuantile | minfiData27k |
| 450k | 482,421 | 135,476 | 346,945 | preprocessRaw, preprocessNoob, preprocessQuantile<br>preprocessFunnorm, preprocessSWAN | minfiData |
| EPIC | 863,904 | 142,262 | 721,642 | preprocessRaw, preprocessNoob, preprocessQuantile<br>preprocessFunnorm, preprocessSWAN | minfiDataEPIC |

### Combining 450k and EPIC arrays by using common probes

Unlike the transition from 27k to 450k arrays, a substantial percentage (93.3%) of loci contained on the 450k microarray are also contained on the EPIC microarray, measured using the exact same probes and same chemistry. This represents a total of 453,093 CpG loci in common between the two arrays. In addition to probes measuring CpG loci, both arrays contain 65 SNP probes to assess possible sample mislabeling, 59 of which are shared by both arrays. Finally, the two arrays both contain a number of control probes, 850 on the 450k array and 635 on the EPIC array, with 487 probes in common.

The large percentage of common loci measured by common probes makes it possible to combine data from 450k and EPIC arrays. The lowest level of the combination can occur at the probe level, before the probes are summarized into methylated and unmethylated intensities. We have implemented this functionality in the function combineArrays which outputs an object that behaves either as a 450k or an EPIC array as chosen by the user with a reduced number of probes; we call this is a virtual array. We achieve this goal by remapping probes between identifiers used on the two arrays. We also support the combination of the two array types at the CpG locus level, after the creation of the methylation and unmethylation channels, with or without prior normalization. This is implemented in the same function.

Most of the common probes between the 27k array and the 450k array (or between the 27k array and the EPIC array) do not have the same probe chemistry, and therefore it is not possible the combine those two arrays at the probe level. Only the creation of a virtual array at the CpG locus level is supported, also via the function combineArrays.

Combining the 450k and EPIC arrays at the probe level has several advantages. First, it opens up the possibility of using advanced preprocessing methods such as Noob (Triche et al., 2013) and functional normalization (Fortin, Labbe, et al., 2014), and using the batch effect correction tools ComBat (Johnson, Li, and Rabinovic, 2007), SVA (Leek and Storey, 2007; Leek and Storey, 2008) and RUVm (Maksimovic, Gagnon-Bartsch, et al., 2015). Second, it also allows users to estimate cell type proportions using existing algorithms (Houseman et al., 2012) making use of flow-sorted reference datasets currently available from whole blood, cord blood and prefrontal cortex assayed on the 450k array (Reinius et al., 2012; Bakulski et al., 2016; Guintivano, Aryee, and Kaminsky, 2013). Finally, this also allows the prediction of A/B compartments for estimating the open/closed state of the chromatin using the methodology described in Fortin and Hansen (2015).

### Single sample normalization with ssNoob

Single sample normalization is of great potential benefit to users, particularly for analyzing large datasets which arrive in batches, because data can be processed separately and independently of the previously processed data. Of particular interest are single sample normalization methods which perform comparable to methods that pool information across multiple samples (McCall, Bolstad, and Irizarry, 2010; Piccolo, Sun, et al., 2012; Piccolo, Withers, et al., 2013). We note that single-sample processing does not remove the need for proper experimental design (Birney, Smith, and Greally, 2016) and, where appropriate, batch effect correction methods such as ComBat (Johnson, Li, and Rabinovic, 2007), SVA (Leek and Storey, 2007; Leek and Storey, 2008) and RUVm (Maksimovic, Gagnon-Bartsch, et al., 2015), especially for large or multi-center studies. If the incrementally processed results are directly comparable to batch-processed results from large studies (such as the Cancer Genome Atlas project, or TCGA), the benefits of single sample processing are magnified.

We determined that the Noob method (Triche et al., 2013), which combines both background correction and dye bias equalization, behaves as a single sample normalization method for Beta values: the Beta values computed after Noob normalization are the same whether samples are processed individually or in a group (see below for a proof). This is not currently the case for the methylated and unmethylated probe intensities themselves. Therefore, we updated the Noob algorithm to remove the need for a reference sample, thereby creating a full single sample normalization method with excellent performance. We emphasize that this change to the Noob method does not affect the resulting Beta values, and ssNoob-processed arrays are directly comparable to Level 3 TCGA methylation array data, as the latter were processed with the reference-based Noob algorithm. The new version of Noob is now the default from minfi version 1.19.10.

For single sample preprocessing, we eliminated the use of a reference sample *r* in correcting the dye bias ratio *R*_*UM*_ estimated from normalization controls (*c*_*AT*_ for *U*, *c*_*GC*_ for *M*). Previously (Triche et al., 2013), we computed the corrected 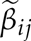 at probe *j* in sample *i* as

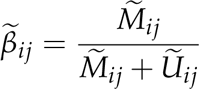

where the corrected methylated and unmethylated intensities are estimated as

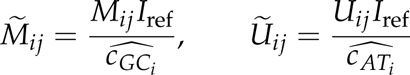

with

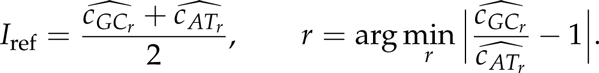

Here the methylated *M*_*ij*_ and unmethylated intensities *U*_*ij*_ are background corrected as described in Triche et al. (2013), and 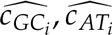 are the averages of the normalization controls on array *i*. We note that background correction is a single sample procedure.

Note that, since the reference normalization control intensity *I*_*ref*_ vanishes when we convert the corrected 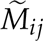 and 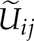 into 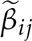, it has no impact on the corrected Beta values. We address the dye bias in Type II probes by correcting *U*_*ij*_ as

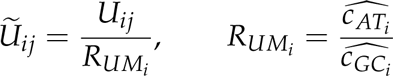

recovering the dye bias corrected Beta values as

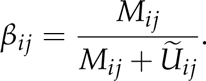

The resulting *β*_*ij*_ is identical to reference-based 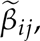, and accuracy of detection p-values (based on intensities) may also benefit from the reduced manipulation of raw intensities.

### Background correction & dye bias correction reduce technical variation

We assessed how the different preprocessing methods available in minfi perform at reducing technical variation among three technical replicates of the cell line GM12878, bundled in the minfiDataEPIC package. The different methods are: preprocessing as Illumina, SWAN normalization (Maksimovic, Gordon, and Oshlack, 2012), stratified quantile normalization as discussed in Touleimat and Tost (2012) and Aryee et al. (2014), single-sample noob background correction followed by dye bias correction (Triche et al., 2013), and functional normalization (Fortin, Labbe, et al., 2014). We also compare those methods to no normalization.

To assess technical variation, we calculated the variance of the Beta values across the the 3 technical replicates at each CpG, stratified by probe design type. Boxplots of the distribution of these variances are shown in Figure 1. The results follow the patterns that we have reported previously (Fortin, Labbe, et al., 2014), and show that the EPIC array behaves similarly to the 450k array on this assessment. This result is limited by the following: (1) it is based on only 3 technical replicates of a single sample and (2) technical variability is a poor proxy for performance, as we have shown previously (Fortin, Labbe, et al., 2014). To expand on the second point, we have shown previously that methods which best reduce technical variation are not the best at achieving replication between different datasets, where the latter is arguably of far greater importance.

**Figure 1.**
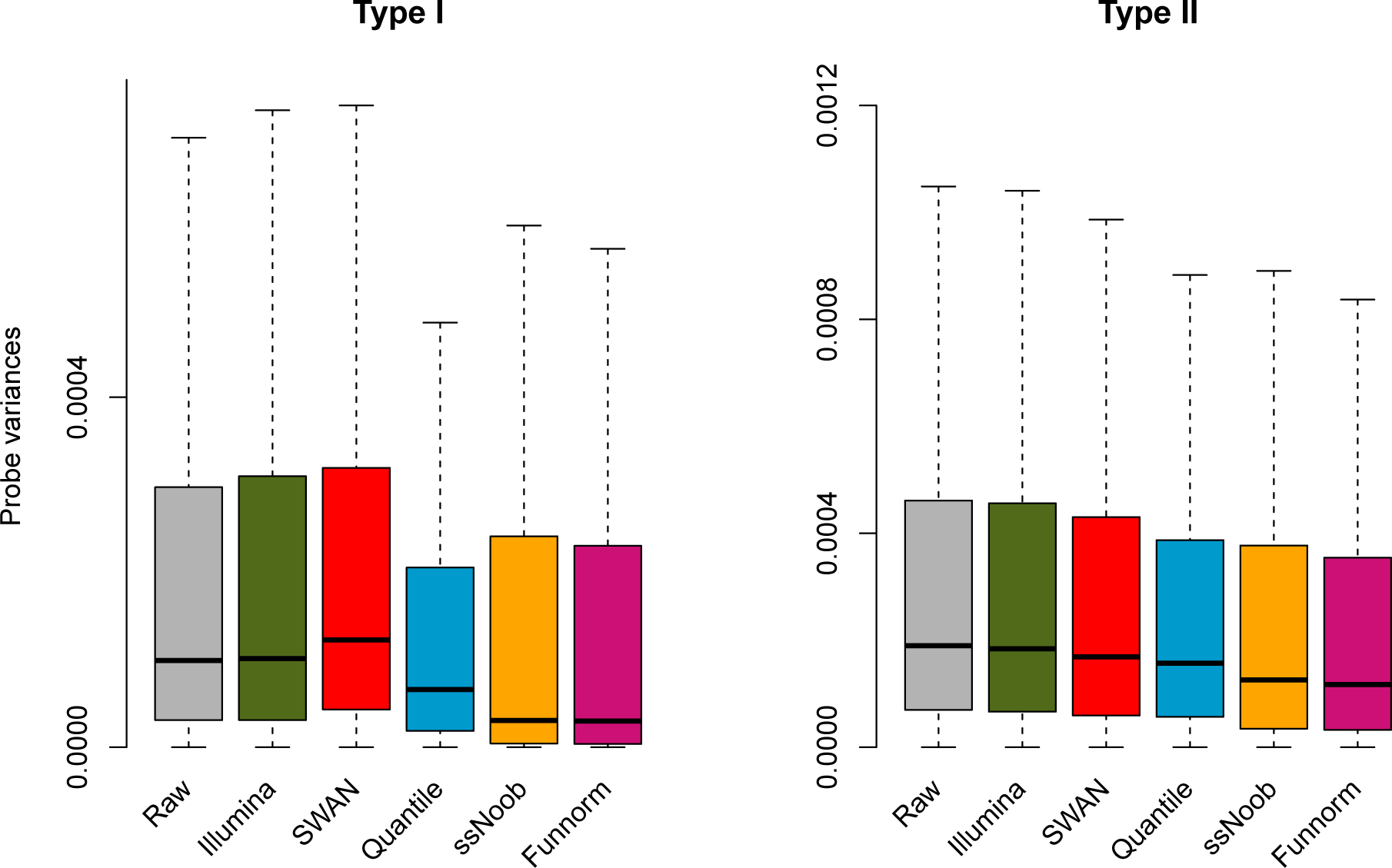
Variances between 3 technical replicates assayed on the EPIC array. Data was normalized using various normalization methods and a variance across the three technical replicates was computed for each methylation loci. We show the distribution of these variances.

### Background correction & dye bias correction improve sample classification across array types

Above, we describe how to combine data from the EPIC and 450k array into a virtual array that contains only common probes or CpG loci. This implies discarding probes or CpG loci only measured on one of the array types. It might be of interest for researchers to normalize the two array types without discarding these probes or CpG loci. Most normalization procedures for the 450k array modify the marginal distributions of each sample by estimating a global trend across all samples. For such procedures, it can have unfortunate results if different subsets of the data are normalized separately. We therefore expected that separate normalization of EPIC data and 450k data might lead to reduced performance.

To examine this question, we used the three technical replicates of the GM12878 cell line assayed on the EPIC array, and compared them to a set of 450k arrays we combined from publicly available data (Table 2). This set consists of 261 lymphoblastoid cell lines (LCLs), the same cell type as GM12878, and 78 other samples. The 78 other samples have a selection of 20 peripheral blood mononuclear (PBMC) samples as well as 58 samples from ENCODE.

First we assessed the performance of the different normalization methods, when the data was combined at the probe level into a virtual array and subsequently normalized together. As performance measure, we computed the median distance between the data from the EPIC array and all of the 450k data. A useful normalization strategy will result in the LCLs drawing closer to each other while moving further from the other cell types. We used the distance as a metric for predicting whether or not a 450k sample is an LCL sample, and displayed prediction performance as an ROC curve (Figure 2a). While all methods predict well, this shows that ssNoob, functional normalization and quantile normalization achieved perfect prediction performance. To investigate whether or not the methods can separate the PBMC samples from the ENCODE samples, we plotted the sorted values of the distance between the EPIC data and the 450k samples (Figure 3), and in Figure 5, we show the median distance between the ENCODE GM12878 cell line assayed on the 450k platform and the EPIC arrays (full dots). ssNoob performs the best, followed by functional normalization and quantile normalization. These assessments broadly mirror existing literature (Fortin, Labbe, et al., 2014).

**Figure 2.**
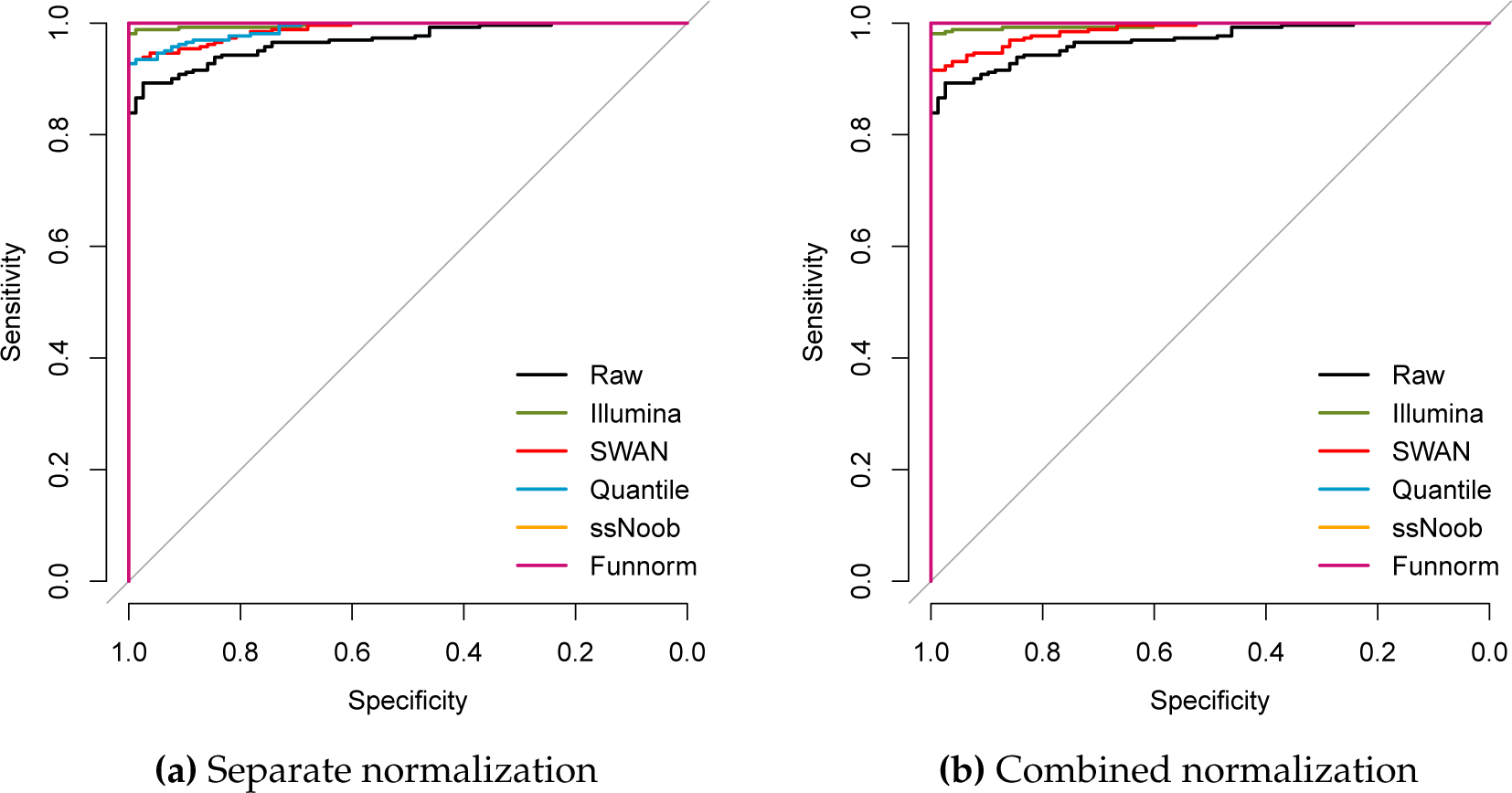
Normalization assessment using ROC curves. The median distance between LCLs measured on the EPIC array and a number of different samples measured using the 450k array was used to predict whether a 450k sample was a LCL or not. Displayed is an ROC curve showing the performance of the predictor. (a) EPIC and 450k samples were combined into a virtual array and then subsequently normalized together. (b) EPIC and 450k samples were normalized separately and then subsequently combined at the methylation loci level.

**Figure 3.**
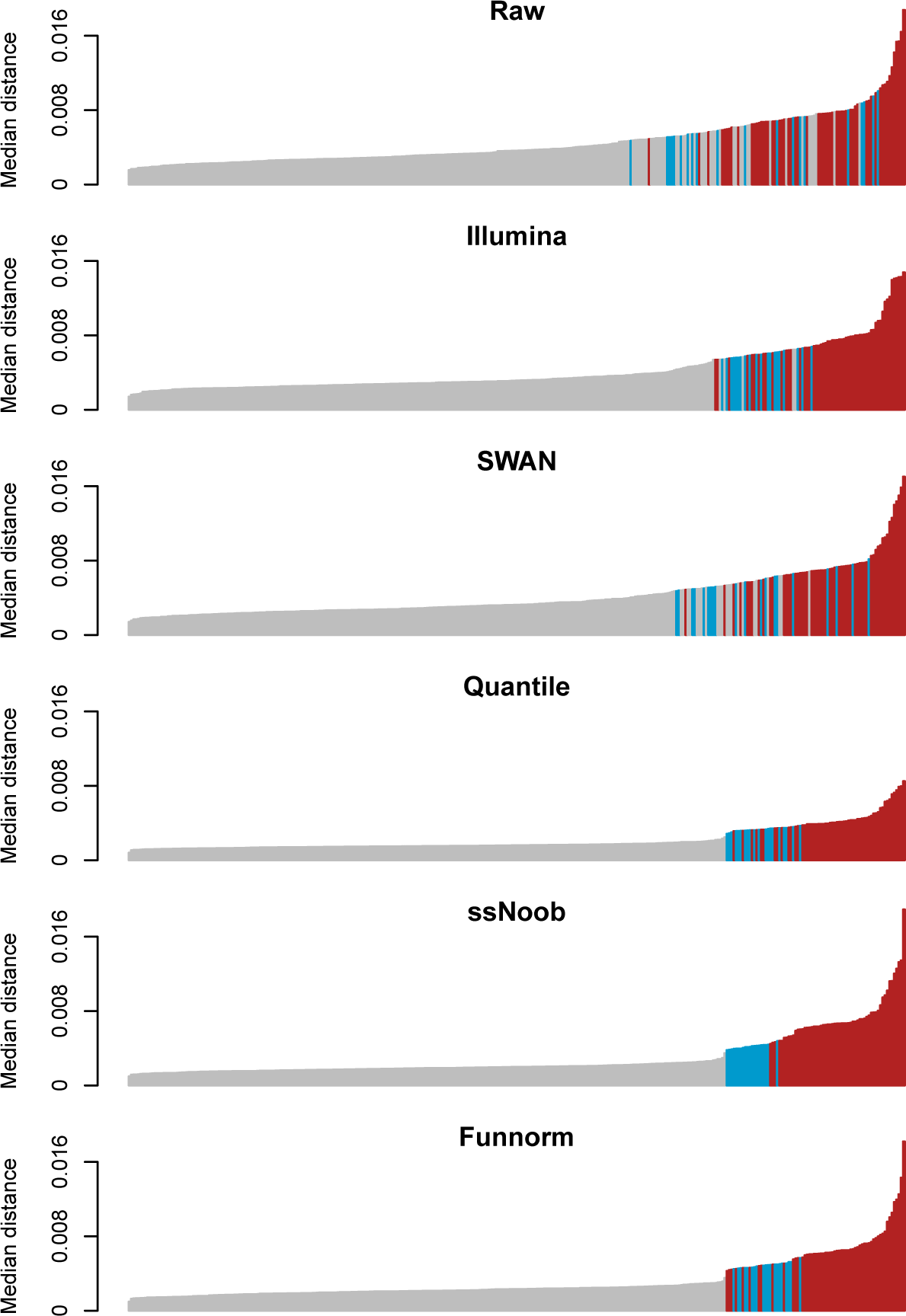
Median distance to EPIC array for normalization on combined virtual array. The median distance between LCLs measured on the EPIC array and a number of different samples measured using the 450k array. LCL samples are depicted in grey (261 in total), PBMC samples are depicted in blue (20 samples) and measurements of various ENCODE cell lines are depicted in blue (58 samples). All samples (both EPIC and 450k) were combined into a virtual array prior to normalization.

Next, we explored the effect of normalizing the 450k samples separately from the EPIC samples followed by combining the data at the CpG loci level. We used the same assessments to investigate performance (Figures 2b, 4, and circle dots in Figure 5). The main difference is that quantile normalization performs significantly worse. This is expected: quantile normalization normalizes data to a common reference which is empirically determined. By separating the EPIC and 450k data, the reference distribution will differ between the two datasets. Given the performance of ssNoob, together with the fact that functional normalization might suffer from the same issue as quantile normalization, we recommend ssNoob for separate normalization of EPIC and 450k arrays, if the goal is to subsequently combine the data from the two arrays. We note that it may be possible to extend functional normalization and quantile normalization to use targeted quantile normalization and ensure that the reference is common across the two array types.

**Figure 4.**
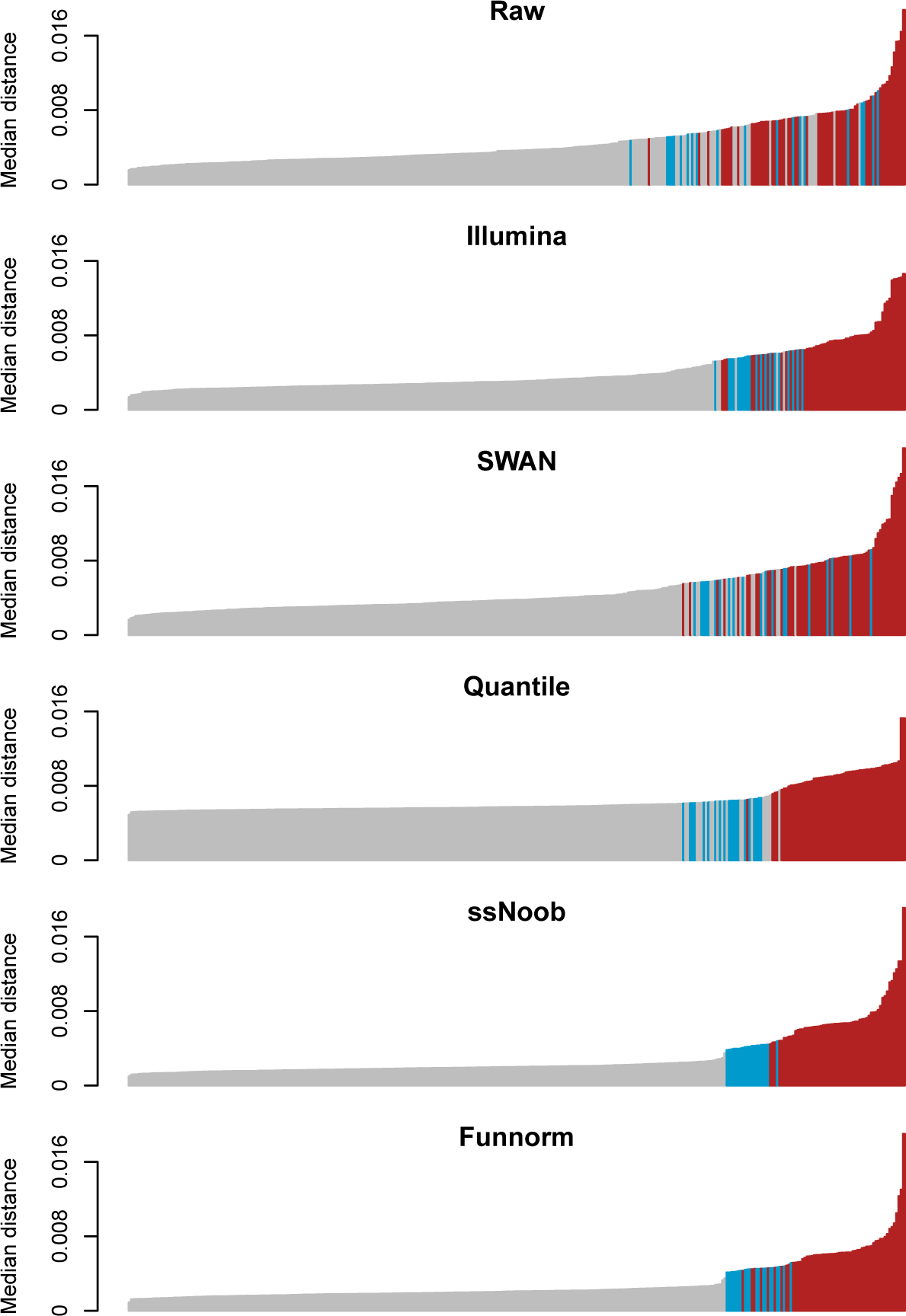
Median distance between 450k samples and the average EPIC array sample for normalization on separate arrays. As Figure 3, but EPIC and 450k data were normalized separately and subsequently combined.

**Figure 5.**
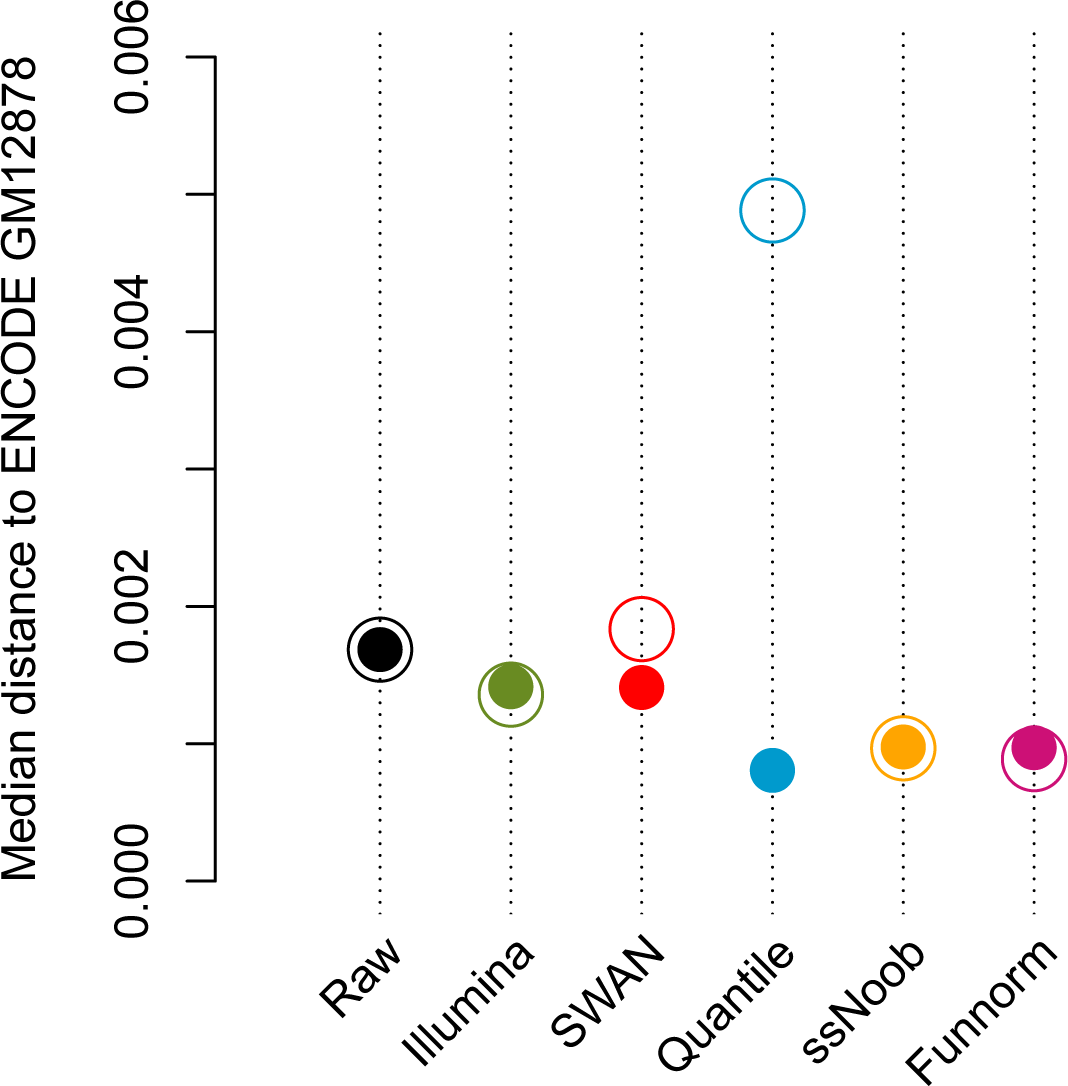
Median distance between the 450k ENCODE GM12878 sample and the average EPIC array sample for different normalizations. The full dots represent the median distances for data normalized after the creation of a virtual array, while the circles represent the median distances when the EPIC and 450k data are normalized separately and subsequently combined.

### Estimation of cell-type composition for EPIC arrays

It has been shown that in epigenome-wide association studies (EWAS), celullar heterogeneity, that is different cell-type proportions across samples, can lead to spurious associations when confounded with the outcome of interest (Houseman et al., 2012; Jaffe and Irizarry, 2014). Several methods have been proposed to estimate the cell-type proportions from a reference dataset that is made of sorted samples (Houseman et al., 2012; Jaffe and Irizarry, 2014). Reference 450k array datasets exist for whole blood, cord blood and prefontal cortex (Reinius et al., 2012; Bakulski et al., 2016; Guintivano, Aryee, and Kaminsky, 2013) and have been made available via the Bioconductor packages “FlowSorted.Blood.450k”, “FlowSorted.CordBlood.450k” and “FlowSorted.DLPFC.450k”, respectively.

The function `estimateCellCounts` in minfi allows users to estimate the cell-type proportions in their dataset using the reference datasets listed above. To take advantage of the existing 450k datasets for the cell-type composition estimation in EPIC arrays, we adapted the `estimateCellCounts` to the EPIC array. Briefly, if an EPIC array dataset is provided, it is first converted to a 450k dataset by removing probes that differ between the two arrays, and then combined with the reference 450k dataset by considering only probes in common. This results in a virtual 450k array dataset that contains 7% less probes than an common 450k dataset.

In order the evaluate how removing 7% of probes from the 450k platform impacts the cell-type composition estimation, we estimated the cell-type proportions for the 20 PBMC samples from the 450k-Esteller dataset, before and after removing the probes that differ between the 450k and EPIC arrays. We estimated the proportions for the cell types CD8T, CD4T, NK, BCell, Mono and Gran using the package `FlowSorted.Blood.450k` as a reference dataset. Both the reference dataset and the 450k-Esteller dataset were normalized together using quantile normalization. We note that other choices of normalization can be applied as well. This yielded very good results; for each cell type, the correlation of the cell type proportions between the full dataset and the probe-reduced dataset is higher than 0.99, with a negligible average difference of 0.001 between the two sets of proportions.

## Discussion

We have described our work to support EPIC and 27k arrays in the widely used minfi package. Our primary contribution has been to develop software which easily handles and combines data from the different arrays. We also evaluated standard normalization procedures and shown that they perform as expected for the EPIC array, based on the literature on the 450k array. And finally, we have modified the noob method to be a true single sample normalization method.

By combining data from the EPIC and the 450k array at the probe level, we have made it possible to jointly process data from the two platforms together, using state of the art methods. In addition, it allows users to estimate cell type proportions for samples assayed on the EPIC array using reference datasets measured on the 450k array.

For normalization, we conclude that the single sample noob method is superior for joint analysis of EPIC and 450k data, followed by functional normalization and quantile normalization. We show that care We show that care must be taken if the two array types are normalized separately and later combined.

Our evaluation of different normalization procedures is limited by the fact that the available EPIC data is 3 technical replicates of an lymphoblastoid cell line. Forthcoming studies with biological specimens, where material from each subject was run on each generation of Infinium array, will provide a basis to evaluate the generality of our findings. However, we stress that our results fit expectations based on our general understanding of normalization procedures, and we therefore expect these results to hold under further evaluations.

## Methods

### Data

Data sources used in this paper are listed in Table 2.

**Table 2.** Methylation datasets.

| Dataset | Cell type | n | Type Replicates | Platform | Accession | Reference |
| --- | --- | --- | --- | --- | --- | --- |
| EPIC-LCL | LCL (GM12878) | 3 | Technical | EPIC | - | Illumina website, also available in minfiDataEPIC (Fortin and Hansen, 2016) |
| 450k-Esteller | PBMC | 20 | Biological | 450k | GSE36369 | Heyn et al. (2013) |
|  | LCL | 256 | Biological | 450k | GSE36369 | Heyn et al. (2013) |
| 450k-ENCODE | LCL (GM12878) | 1 | - | 450k | GSE40699 | ENCODE Project Consortium (2004) |
|  | LCL (Others) | 4 | Biological | 450k | GSE40699 | ENCODE Project Consortium (2004) |
|  | Others | 58 | Biological | 450k | GSE40699 | ENCODE Project Consortium (2004) |

### Normalization Assessment

#### Normalization on separate arrays

For each normalization method, we normalized the EPIC and 450k data separately. We then combined the two normalized array datasets at the CpG level, resulting in a matrix of Beta values **B**^Sep^ with 453,093 rows and 342 columns (loci only measured on the EPIC was discarded for our assessment).

#### Normalization on virtual array

We combined the unnormalized EPIC and 450k data at the probe level to create a virtual array. We then normalized the data jointly by applying each normalization method to the combined virtual array, resulting in a matrix of Beta values **B**^Comb^ with 453,093 rows and 342 columns.

#### Median distance of the 450k samples relative to the EPIC array

For both **B**^Sep^ and **B**^Comb^, we compute the average methylation Beta value profile of the GM12878 cell line assayed on the EPIC array by taking the mean of the three technical replicates, resulting in the two vectors of Beta values 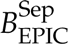 and 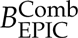. For each normalized 450k sample 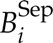 and 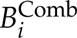, we calculate the median distances 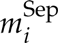 and 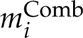 with respect to the EPIC array as follows:

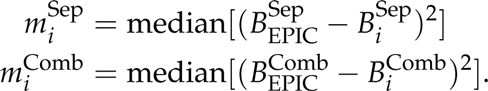

The median distance values 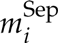 and 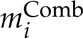 quantify the similarity of each 450k sample with respect to the cell line GM12878 sample assayed on the EPIC array. We present the ordered values 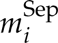 and 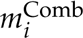 in Figures 4 and 3 respectively. These barplots are colored by tissue type.

For each normalization method, we use the median distances as predictor of cell type (LCL or not LCL) for the 450k samples, and use an ROC curve to summarize the specificity and sensitivity of each normalization method (Figure 2).

### Software

The results in this manuscript were produced using minfi version 1.19.10, minfiDataEPIC version 0.99.3, IlluminaHumanMethylationEPICmanifest version 0.3.0, and IlluminaHumanMethylationEPICanno.ilm10b2.hg19 version 0.3.0. Scripts describing our reproducible analysis are available at https://github.com/hansenlab/EPIC450k_repro.

## Acknowledgements

### Funding

Research reported in this publication was supported by the National Cancer Institute of the National Institutes of Health under award number U24CA180996. Dr. Triche gratefully acknowledges support from the Leukemia & Lymphoma Society Quest for Cures program, the Tower Cancer Research Foundation, the St. Baldrick’s Foundation Pathway Directed Treatment for Refractory AML Consortium grant, and the Jane Anne Nohl Hematology Research Support Fund.

### Disclaimer

The content is solely the responsibility of the authors and does not necessarily represent the official views of the National Institutes of Health.

### Conflict of Interest

None declared.

## Bibliography

Aryee, M. J., A. E. Jaffe, H. Corrada-Bravo, C. Ladd-Acosta, A. P. Feinberg, K. D. Hansen, and R. A. Irizarry (2014). “Minfi: a flexible and comprehensive Bioconductor package for the analysis of Infinium DNA methylation microarrays”. Bioinformatics 30.10, pp. 1363–1369. DOI: 10.1093/bioinformatics/btu049.

Bakulski, K. M., J. I. Feinberg, S. V. Andrews, J. Yang, S. Brown, S.L McKenney, F. Witter, J. Walston, A. P. Feinberg, and M. D. Fallin (2016). “DNA methylation of cord blood cell types: Applications for mixed cell birth studies”. Epigenetics 11.5, pp. 354–362. DOI: 10.1080/15592294.2016.1161875.

Bibikova, M., B. Barnes, C. Tsan, V. Ho, B. Klotzle, J. M. Le, D. Delano, L. Zhang, G. P. Schroth, K. L. Gunderson, J.-B. Fan, and R. Shen (2011). “High density DNA methylation array with single CpG site resolution”. Genomics 98.4, pp. 288–295. DOI: 10.1016/j.ygeno.2011.07.007.

Birney, E., G. D. Smith, and J. M. Greally (2016). “Epigenome-wide Association Studies and the Interpretation of Disease -Omics”. PLoS Genetics 12.6, e1006105. DOI: 10.1371/journal.pgen.1006105.

Cancer Genome Atlas Research Network (2013). “Comprehensive molecular characterization of clear cell renal cell carcinoma”. Nature 499.7456, pp. 43–49. DOI: 10.1038/nature12222.

ENCODE Project Consortium (2004). “The ENCODE (ENCyclopedia Of DNA Elements) Project”. Science 306.5696, pp. 636–640. DOI: 10.1126/science.1105136.

Fortin, J.-P., E. Fertig, and K. Hansen (2014). “shinyMethyl: interactive quality control of Illumina 450k DNA methylation arrays in R”. F1000Research 3, p. 175. DOI: 10.12688/f1000research.4680.2.

Fortin, J.-P. and K. D. Hansen (2015). “Reconstructing A/B compartments as revealed by Hi-C using long-range correlations in epigenetic data”. Genome Biology 16, p. 180. DOI: 10.1186/s13059–015–0741-y.

Fortin, J.-P. and K. D. Hansen (2016). minfiDataEPIC: Example data for the Illumina Methylation EPIC array. R package version 0.99.3. URL: http://www.bioconductor.org/packages/minfiDataEPIC.

Fortin, J.-P., A. Labbe, M. Lemire, B. W. Zanke, T. J. Hudson, E. J. Fertig, C. M. Greenwood, and K. D. Hansen (2014). “Functional normalization of 450k methylation array data improves replication in large cancer studies”. Genome Biology 15.12, p. 503. DOI: 10.1186/s13059–014–0503–2.

Gentleman, R. C., V. J. Carey, D. M. Bates, B. Bolstad, M. Dettling, S. Dudoit, B. Ellis, L. Gautier, Y. Ge, J. Gentry, K. Hornik, T. Hothorn, W. Huber, S. Iacus, R. Irizarry, F. Leisch, C. Li, M. Maechler, A. J. Rossini, G. Sawitzki, C. Smith, G. Smyth, L. Tierney, J. Y. H. Yang, and J. Zhang (2004). “Bioconductor: open software development for computational biology and bioinformatics”. Genome Biology 5.10, R80. DOI: 10.1186/gb-2004–5–10-r80.

Guintivano, J., M. J. Aryee, and Z. A. Kaminsky (2013). “A cell epigenotype specific model for the correction of brain cellular heterogeneity bias and its application to age, brain region and major depression”. Epigenetics 8.3, pp. 290–302. DOI: 10.4161/epi.23924.

Heyn, H., S. Moran, I. Hernando-Herraez, S. Sayols, A. Gomez, J. Sandoval, D. Monk, K. Hata, T. Marques-Bonet, L. Wang, and M. Esteller (2013). “DNA methylation contributes to natural human variation.” Genome Research 23.9, pp. 1363–1372. DOI: 10.1101/gr.154187.112.

Hicks, S. C. and R. A. Irizarry (2015). “quantro: a data-driven approach to guide the choice of an appropriate normalization method”. Genome Biology 16, p. 117. DOI: 10.1186/ s13059–015–0679–0.

Houseman, E. A., W. P. Accomando, D. C. Koestler, B. C. Christensen, C. J. Marsit, H. H. Nelson, J. K. Wiencke, and K. T. Kelsey (2012). “DNA methylation arrays as surrogate measures of cell mixture distribution”. BMC Bioinformatics 13, p. 86. DOI: 10.1186/1471–2105–13–86.

Hovestadt, V. and M. Zapatka (2016). conumee: Enhanced copy-number variation analysis using Illumina 450k methylation arrays. R package version 0.99.4. URL: http://www.bioconductor.org/packages/conumee.

Huber, W., V. J. Carey, R. Gentleman, S. Anders, M. Carlson, B. S. Carvalho, H. C. Bravo, S. Davis, L. Gatto, T. Girke, R. Gottardo, F. Hahne, K. D. Hansen, R. A. Irizarry, M. Lawrence, M. I. Love, J. MacDonald, V. Obenchain, A. K. Oleś, H. Pagès, A. Reyes, P. Shannon, G. K. Smyth, D. Tenenbaum, L. Waldron, and M. Morgan (2015). “Orchestrating high-throughput genomic analysis with Bioconductor”. Nature Methods 12.2, pp. 115–121. DOI: 10.1038/nmeth.3252.

Iterson, M. van, E. W. Tobi, R. C. Slieker, W. den Hollander, R. Luijk, P. E. Slagboom, and B. T. Heijmans (2014). “MethylAid: visual and interactive quality control of large Illumina 450k datasets”. Bioinformatics 30.23, pp. 3435–3437. DOI: 10.1093/bioinformatics/btu566.

Jaffe, A. E. and R. A. Irizarry (2014). “Accounting for cellular heterogeneity is critical in epigenome-wide association studies”. Genome biology 15.2, R31. DOI: 10.1186/gb-2014–15–2-r31.

Johnson, W. E., C. Li, and A. Rabinovic (2007). “Adjusting batch effects in microarray expression data using empirical Bayes methods.” Biostatistics 8.1, pp. 118–127. DOI: 10.1093/biostatistics/kxj037.

Leek, J. T. and J. D. Storey (2007). “Capturing heterogeneity in gene expression studies by surrogate variable analysis”. PLoS Genetics 3.9, pp. 1724–1735. DOI: 10.1371/ journal.pgen.0030161.

– (2008). “A general framework for multiple testing dependence”. Proceedings of the National Academy of Sciences 105.48, pp. 18718–18723. DOI: 10.1073/pnas.0808709105.

Maksimovic, J., J. A. Gagnon-Bartsch, T. P. Speed, and A. Oshlack (2015). “Removing unwanted variation in a differential methylation analysis of Illumina HumanMethylation450 array data”. Nucleic Acids Research 43.16, e106–e106.

Maksimovic, J., L. Gordon, and A. Oshlack (2012). “SWAN: Subset quantile Within-Array Normalization for Illumina Infinium HumanMethylation450 BeadChips.” Genome Biology 13.6, R44. DOI: 10.1186/gb-2012–13–6-r44.

McCall, M. N., B. M. Bolstad, and R. A. Irizarry (2010). “Frozen robust multiarray analysis (fRMA)”. Biostatistics 11.2, pp. 242–253. DOI: 10.1093/biostatistics/kxp059.

Moran, S., C. Arribas, and M. Esteller (2016). “Validation of a DNA methylation microarray for 850,000 CpG sites of the human genome enriched in enhancer sequences”. Epigenomics 8.3, pp. 389–399. DOI: 10.2217/epi.15.114.

Morris, T. J., L. M. Butcher, A. Feber, A. E. Teschendorff, A. R. Chakravarthy, T. K. Wojdacz, and S. Beck (2014). “ChAMP: 450k Chip Analysis Methylation Pipeline”. Bioinformatics 30.3, pp. 428–430. DOI: 10.1093/bioinformatics/btt684.

Oros Klein, K., S. Grinek, S. Bernatsky, L. Bouchard, A. Ciampi, I. Colmegna, J.-P. Fortin, L. Gao, M.-F. Hivert, M. Hudson, M. S. Kobor, A. Labbe, J. L. MacIsaac, M. J. Meaney, A. M. Morin, K. J. O’Donnell, T. Pastinen, M. H. Van Ijzendoorn, G. Voisin, and C. M. T. Greenwood (2016). “funtooNorm: an R package for normalization of DNA methylation data when there are multiple cell or tissue types”. Bioinformatics 32.4, pp. 593–595. DOI: 10.1093/bioinformatics/btv615.

Papillon-Cavanagh, S., J.-P. Fortin, and N. De Jay (2013). CopyNumber450k: R package for calling CNV from Illumina 450k methylation microarrays. R package version 1.9.0. URL: http://www.bioconductor.org/packages/CopyNumber450k.

Peters, T. J., M. J. Buckley, A. L. Statham, R. Pidsley, K. Samaras, R. V Lord, S. J. Clark, and P. L. Molloy (2015). “De novo identification of differentially methylated regions in the human genome”. Epigenetics & Chromatin 8, p. 6. DOI: 10.1186/1756–8935–8–6.

Phipson, B., J. Maksimovic, and A. Oshlack (2016). “missMethyl: an R package for analyzing data from Illumina’s HumanMethylation450 platform”. Bioinformatics 32.2, pp. 286–288. DOI: 10.1093/bioinformatics/btv560.

Piccolo, S. R., Y. Sun, J. D. Campbell, M. E. Lenburg, A. H. Bild, and W. E. Johnson (2012). “A single-sample microarray normalization method to facilitate personalized-medicine workflows”. Genomics 100.6, pp. 337–344. DOI: 10.1016/j. ygeno.2012.08.003.

Piccolo, S. R., M. R. Withers, O. E. Francis, A. H. Bild, and W. E. Johnson (2013). “Multiplatform single-sample estimates of transcriptional activation”. Proceedings of the National Academy of Sciences of the United States of America 110.44, pp. 17778–17783. DOI: 10.1073/pnas.1305823110.

Reinius, L. E., N. Acevedo, M. Joerink, G. Pershagen, S.-E. Dahlén, D. Greco, C. Söderhäall, A. Scheynius, and J. Kere (2012). “Differential DNA methylation in purified human blood cells: implications for cell lineage and studies on disease susceptibility”. PLOS ONE 7.7, e41361. DOI: 10.1371/journal.pone.0041361.

Ruiz-Arenas, C., C. Hernandez-Ferrer, and J. R. Gonzalez (2016). MultiDataSet: Implementation of the BRGE’s (Bioinformatic Research Group in Epidemiology from Center for Research in Environmental Epidemiology) MultiDataSet and MethylationSet. R package version 1.1.2. URL: http://www.bioconductor.org/packages/MultiDataSet.

Touleimat, N. and J. Tost (2012). “Complete pipeline for Infinium Human Methylation 450K BeadChip data processing using subset quantile normalization for accurate DNA methylation estimation.” Epigenomics 4.3, pp. 325–341. DOI: 10.2217/epi.12.21.

Triche, T. J., D. J. Weisenberger, D. Van Den Berg, P. W. Laird, and K. D. Siegmund (2013). “Low-level processing of Illumina Infinium DNA Methylation BeadArrays”. Nucleic Acids Research 41.7, e90. DOI: 10.1093/nar/gkt090.

Xu, Z., L. Niu, L. Li, and J. A. Taylor (2016). “ENmix: a novel background correction method for Illumina HumanMethylation450 BeadChip”. Nucleic Acids Research 44.3, e20. DOI: 10.1093/nar/gkv907.

Yao, L., H. Shen, P. W. Laird, P. J. Farnham, and B. P. Berman (2015). “Inferring regulatory element landscapes and transcription factor networks from cancer methylomes”. Genome Biology 16, p. 105. DOI: 10.1186/s13059–015–0668–3.

